# Nutritive diet becomes unhealthy if taken in a wrong ratio: Scientific validation of Ayurvedic concept of “incompatible diet” (*Virrudh Aahar*)

**DOI:** 10.1101/708859

**Authors:** Prerana Aditi, Shivani Srivastva, Harsh pandey, Y.B. Tripathi

## Abstract

Honey and ghee is an important constituent of our diet. Honey contain about 200 components in its which play an important role as antioxidative, anti-inflammatory, antimicrobial, wound healing and sunburn, etc. Ghee has also many essential constituents such as phospholipids, PUFA, fat- soluble vitamins, etc. It is mention in Charak Samhita that equal ratio of honey and ghee when taken in food become toxic to health. This study was designed to explore the mechanism of toxicity through the biochemical and histological parameter. For this study, 24 Charles foster rats were taken and divided into four groups (n=6) normal, honey, ghee, and honey+ghee (1:1). After 60 days rats were sacrificed. Weight loss, Hair loss and red patches on ear were found on honey+ghee fed rats. LFT, amadori test, AGE, UV, glucose, DPP-4 and LPO were found significantly increased in honey+ghee group. SOD, Catalase, GSH, ABTS+, GLP-1, GIP-1, ACB and HB were found significantly decreased in honey+ghee group. No changes were found in RFT. H&E show normal morphology in kidney, intestine, and pancreas but mild inflammation was found in liver tissue. We conclude that the equal mixture of honey and ghee shows toxic effect. Increased formation of hydroxymethyl furfural, amadori product, DPP-4 activity and low incretins (GLP-1, GIP-1) activity resulting high postprandial hyperglycemic response are collectively responsible for oxidative stress-mediated toxicity of the mixture of honey and ghee in equal ratio.

## 1. Introduction

Honey and ghee are important constituents of the normal diet. Honey is a sweet, thick, yellowish liquid made by honey bees mainly by sugar-rich nectar of the flower. It is a complex mixture having more than 200 substances(Escuredo, Míguez, Fernández-González, & Carmen Seijo, 2013). These are in consistence because of variation in type of flower nectar which is used by the honey bee. Rather than this floral source, some external factor is also responsible for the variability in honey composition such as environmental and seasonal factor and processing time or way, time of storage and packing (Escuredo, Dobre, Fernández-González, & Seijo, 2014). Honey is a supersaturated solution of sugars, of which fructose (38%) and glucose (31%) is the important contributor. Some minor constituent are also present in wide range which include phenolic acid and flavonoids (Andrade, Ferreres, & Amaral, 1997), certain enzymes like glucose oxidase, catalase, ascorbic acid(Andrade et al., 1997), carotenoid like compound (Tan, Wilkins, Holland, & McGhie, 1989), organic acids (Tan et al., 1989), maillard reaction product, amino acids and protein (Tan et al., 1989), minerals, vitamins ions (Alqarni, Owayss, Mahmoud, & Hannan, 2014). These phenolic compound and flavonoids present in honey is a rich source of antioxidant. Honey is a sweet and flavorful natural product, which is consumed for its high nutritive value and for its effects on human health, with antioxidant, bacteriostatic, anti-inflammatory and antimicrobial properties, as well as wound and sunburn healing effects(Tan et al., 1989).

Ghee is similar to clarified butter fat which is produced by the heating butter to remove milk solids and water. It is prepared from cow’s milk, buffalo milk’s and mixed milk’s. Ghee contain fats (99%), vitamins, cholesterol, tochopherol, free fatty acids, carotene, lanosterol, ubiquinones, etc(Sserunjogi, Abrahamsen, & Narvhus, 1998). Ghee is made from butter but the milk solids and impurities have been removed, so most people who are lactose or casein intolerant have no issue with ghee.

Although both are rich nutrients but as per ayurvedic literature its regular use in equal quantity, in one food, proof be toxic. (Charak Samhita, Charaka Sutra sthana, ½, Page-4, 2001). It is mention in Charak Samhita that when we take this incompatible diet for a short time then it may not be dangerous for health but when we take it regularly for longer time it become dangerous and may be cause types of diseases like sterility, herpes, eye diseases, skin eruptions, ascites, fistula, leprosy, sprue, edema, fever, rhinitis, fetal distress, and may even cause death (Charak Samhita, Charak Sutra sthana, 26/102, page-361, 2001). However this believes has not been scientifically validated till date, though some research papers are available (K. R. Anilakumar, Annapoorani, Murthy, & Bawa, 2011)(K. Anilakumar, Khanum, Murthy, Bawa, & Annapoorani, 2010).

Here we have tried to explore the effect of this mixture on the physical appearance, anatomical and physiological changes on GIT with special reference to oxidative stress, incretin secretion, structural modification in blood component circulating protein and carbohydrates.

## 2. Method and materials

### 2.1. Material

Honey was purchased from Patanjali. Ghee was purchased from Anik. All the biochemical kits were purchased from Accurex Biomedical PVT LTD., Biosar, Thane. Lipase substrate, 4-nitrophenyl butyrate, DPP-4 substrate gly-pro-p-nitroanilide, and ABTS^+^ tablet were purchased from Sigma-Aldrich. GLP-1 and GIP-1 EIA kit was purchased from Sigma-Aldrich. Nitroblue tetrazolium chloride, Riboflavin, Methionine, Bradford reagent, Bromocresol green (BCG) reagent, Thiobarbutiric acid, Tricholoro acetic acid were purchased from Hi-media Ltd, Calcutta.

### 2.2. Method

The experimental protocol was approved by an animal welfare ethic committee of the Institute of Medical Sciences, Banaras Hindu University, Varanasi (ethical committee letter # No Dean/2017/CAEC/720). Healthy male Charles Foster strain rats (24) from an inbreed colony were randomly selected from the central animal house of our institute (Institute of Medical Sciences). They were of average weight ranging between 100–200 gm. The purpose of taking male rat is only to avoid the variation like different metabolic rate of gender including sex hormones, lactation, pregnancy which is inherent in the female. Rats were kept separately in propylene cage, fed standard diet and kept in hygiene conditions. The entire animals were kept in 25° C with standard condition 12 h light/dark cycle. Before treatments start rats were kept under the standard condition for 1 week with free access to standard chow and tap water for acclimatization.

### 2.3. Treatment

After acclimatization, the rats were divided into four groups (n=6). Group-1 received the normal diet. Group-2 received honey along with the normal diet. Group-3 received ghee along with the normal diet. Group-4 received equal mixture of honey and ghee (Honey+ghee) along with the normal diet. For finalization of dose, we have taken the basis of earlier publications where 2.5g/kg/day dose of honey have been used(Sadeghi, Salehi, Kohanmoo, & Akhlaghi, 2019). As per Ayurvedic practice, the recommended dose of honey is 48g/day and the recommended dose of ghee is 30ml/day for a healthy 70 kg men. Based on these literatures we have selected the dose. Honey and ghee were given 0.7 ml/100gm body weight for 60 days. Equal ratio of honey and ghee were mixed and given to rats for 60 days. Just before given, the mixture was vigorously vortex for emulsion the mixture and immediately given by the oral gavages. These experiments were carried out for 60 days and weight, physical appearance, food intake was recorded and blood was collected on every 15 days. Rats were sacrificed on 61 day by using ethyl ether and Liver, kidney, intestine, pancreas organ were collected and washed with 1x PBS for various biochemical and histological study.

### 2.4. Biochemical study

#### 2.4.1. Liver function test

SGOT, SGPT, ALP test were done by using the commercial kit available (Accurex Biomedical). Albumin level was estimated by BCG reagent (Hi-media). Protein estimation was done by Bradford’s reagent (Hi-media).

#### 2.4.2. Kidney function test

Urea, BUN, and Creatinine test were done by using commercial kit available (Accurex Biomedical).

#### 2.4.3. Oxidative stress parameter

Superoxide dismutase (SOD) activity was measured in term of inhibition of reduction of nitro blue tetrazolium (NBT) in the presence of riboflavin as described (Beauchamp & Fridovich, 1971) with slight modification (Nagwani & Tripathi, 2010). Catalase enzyme activity was measured by Aebi’s method by monitoring H_2_O_2_ breakdown at 240 nm (Aebi, 1984). GSH was done according to described by Beutler et.al,(Beutler, Duron, & Kelly, 1963) with slight modification (Tripathi, Shukla, Chaurasia, & Chaturvedi, 1996). ABTS^.+^ radical scavenging activity was determined according to described by Re.et.al. (Re et al.,1999). Lipid peroxidation was measured by the thaiobarbutiric acid method (Matsunami, Sato, Sato, Yukawa, & Sato, 2010).

#### 2.4.4. Protein modification parameter

##### 2.4.4.1. Amadori test

The formation of Amadori products was assessed using the method of Johnson and Baker (Johnson & Baker, 1987), with nitroblue tetrazolium (NBT). One hundred microliters of the sample per well were transferred to a 96-well plate. One hundred microliters of NBT reagent (250 μM NBT in 0.1 M carbonate buffer, pH 10.35) were added to each well, and the plate was incubated at 37 °C for 2 h. Absorbance was measured at 525 nm. An absorption coefficient of 12,640 cm-1 mol-1 for monoformazan was used (Mironova, Niwa, Handzhiyski, Sredovska, & Ivanov, 2005).

##### 2.4.4.2. UV Absorbance Spectroscopy

Absorption profiles of normal and treated plasma were recorded on Perkin Elmer Lambda 25 UV/VIS spectrometer in 200 to 400 nm wavelength range using quartz cuvette of 1 cm path length. Increase in absorbance at 280 nm indicated the increased in hyperchromatocity due to protein oxidation.

##### 2.4.4.3. Fluorescence Study for AGEs

AGE-specific fluorescence was recorded by exciting the samples at 390 nm and keeping the emission range of 400 to 600 nm in PerkinElmer LS 45 fluorescence spectrometer (Jairajpuri, Fatima, & Saleemuddin, 2007).

##### 2.4.4.4. Albumin –cobalt binding assay

The 100□μL of serum was mixed with 25□μL cobalt chloride (CoCl_2_, 1□mg/mL) in a 96-well plate and incubated at room temperature for 10 minutes. The 25□μL dithiothreitol (DTT, 1.5□mg/mL) was added, followed by 2-minute incubation to allow the reaction with free cobalt salt. Finally, 150□μL saline was added to terminate the reaction. Absorbance was measured at 470□nm(Bar-Or, Lau, & Winkler, 2000). % decreased was calculated to show the albumin to cobalt binding.

#### 2.4.5. Glycemic parameter

Glucose level was estimated by using the Accurex Biomedical kit. GLP-1 and GIP-1 enzyme activity was done by using EIA kit (Sigma-Aldrich). DPP-4(dipeptidyl peptidase) activity was done by adding 95 μl GPPN (0.2 mM) as a substrate in the mixture of 65 μl Tris-HCl (50 mM, 7.5 pH) and 10 μl plasma. Absorbance was taken immediately and after 20 min at 405 nm.

Enzyme unit (U/ml) = (((ΔAbs/min (sample)- ΔAbs/min(blank))* reaction volume*dilution factor)/molar extinction coefficient * path length*sample volume

#### 2.4.6. Triglycerides and Lipase activity in plasma

Triglycerides level was estimated by Accurex Biomedical kit. Lipase activity was done according to Winkler and Stuckmann (Winkler & Stuckmann, 1979). Briefly, the reaction mixture was prepared by mixing one volume of a solution containing 10mM of PNPB into isopropanol with 18 part of the solution of phosphate buffer 100mM, pH-7, and 7.0% ±0.5% w/v triton x100. Ten μl of enzyme was added in appropriate dilution and 190 μl of the reaction mixture was placed in ELISA plate and read absorbance at 415nm for 10min and time interval 30s at 30□C.

Enzyme unit (U/ml) = (((ΔAbs/min (sample) - ΔAbs/min (blank))* reaction volume*dilution factor)/molar extinction coefficient * sample volume

### 2.5. Histology study

H& E staining was done in formalin-fixed liver, kidney, pancreas and intestine (jejunum) tissue followed by rehydration, dehydration, block preparation, slide preparation, staining and mounting described in the early procedure (Suvarna, Layton, & Bancroft, n.d.). Image was examined under a microscope (Eclipse 50i; Nikon, Kanagawa, Japan) encumbered with imaging software (NIS Elements Basic Research; Nikon).

### 2.6. Statistical study

Statistical analysis was carried out using one way ANOVA test followed by Post Hoc analysis with Tukey’s test by using SPSS software. All the result were expressed in mean±SD. The statistical significant difference was considered as p-value less than or equal to 0.05.

## 3. Results

### 3.1. Effect on weight and Physical appearances

The weight of normal group rats was 52% increased, in honey group rats weight was 35.2 % increased, in ghee group rats weight was 39.47 % increased on 60 day comparison to 1^st^ day. We found that the weight of the honey+ghee group rats was firstly increased 16.59% till 30 day. After 30 days weight was found to decrease in the honey+ghee group. On 60 day weight was −0.78 % decreased in Honey+Ghee group rats comparison to 1^st^ day (Table: 1). We found normal skin in all 3 groups until 60 days. But in equal ratio of honey and ghee fed rats (honey+ghee) we found the hair loss on 60 day (Figure: 1). We also found the red patches on the ear on honey+ghee group rats on 60 days but there were no any red patches found on normal, honey and ghee group rats (Figure: 2). The result of physical appearance on different days was shown in Table: 2.

**Figure 1:**
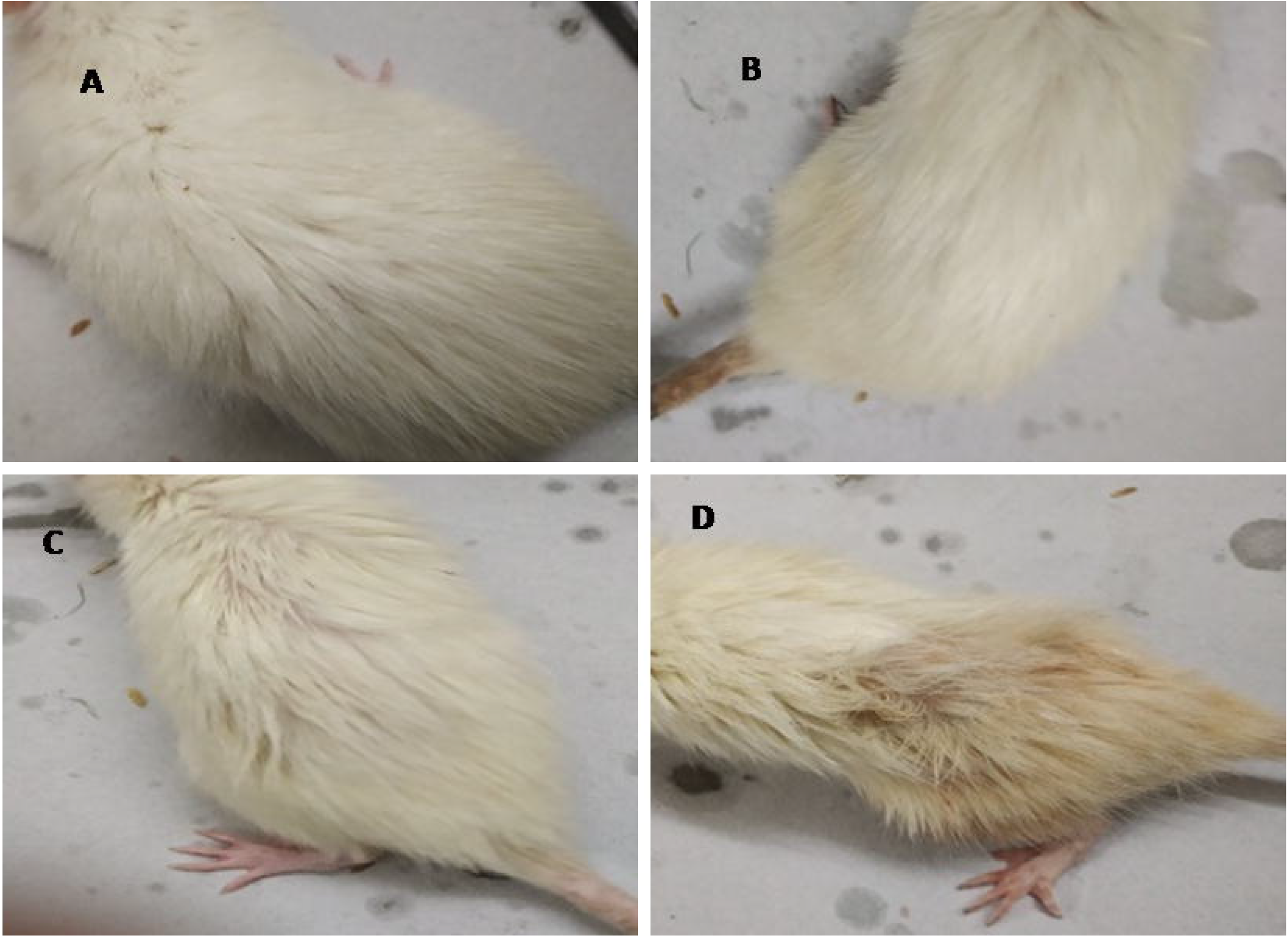
Hair loss and yellowish on skin of rats. A (Normal), B (Honey), C (Ghee), D (Honey+Ghee). Honey + Ghee group showed some loss of hair and yellowish on back skin.

**Figure 2:**
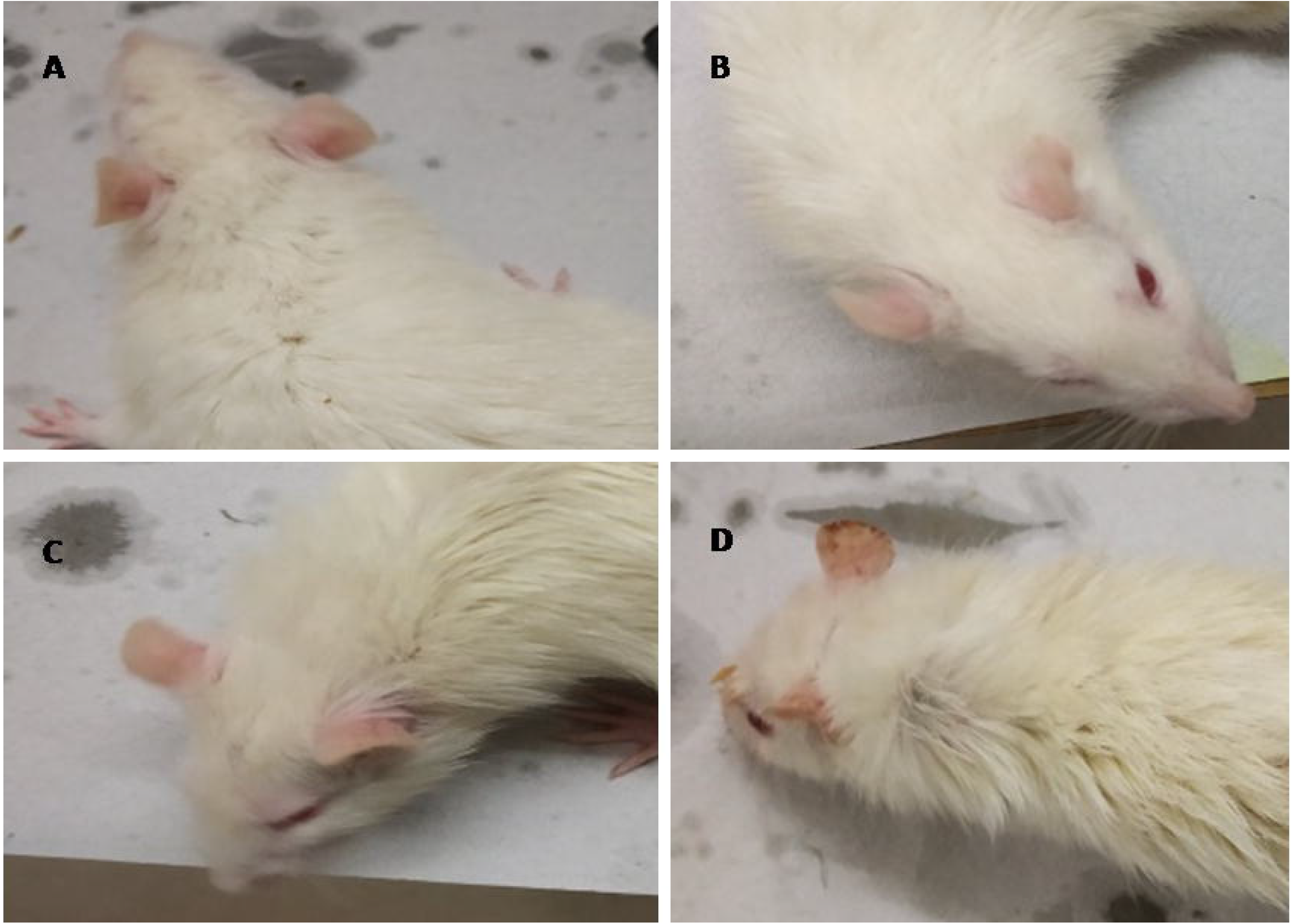
Appearance of red patches on ear of rats. A (Normal), B (Honey), C (Ghee), D (Honey+Ghee). Honey + Ghee group showed appearance of red patches on the ear.

**Table 1:** Weight (gm) of all groups rat on different days

| Days | Normal |  | Honey |  | Ghee |  | Honey+Ghee |  |
| --- | --- | --- | --- | --- | --- | --- | --- | --- |
|  | Weight | % change | Weight | % change | Weight | % change | Weight | % change |
| **1 day** | 120±17.3 | --- | 158±17.5 | --- | 115±15 | --- | 161±27 | --- |
| **15 day** | 130±20.3 | 7.69 | 179±10.3 | 11.18 | 136±25 | 15.85 | 166±24 | 3.6 |
| **30 day** | 180±28.4 | 33.33 | 206±11.5 | 23.15 | 150±14 | 23.33 | 193±23 | 16.59 |
| **45 day** | 210±25.3 | 42.85 | 230±22.5 | 30.97 | 185±21 | 37.83 | 178±25 | 9.58 |
| **60 day** | 250±30.4 | 52 | 245±27.8 | 35.2 | 190±28 | 39.47 | 160±20 | -0.78 |

% change was calculated between 1<sup>st</sup> day and present day.

**Table 2:**
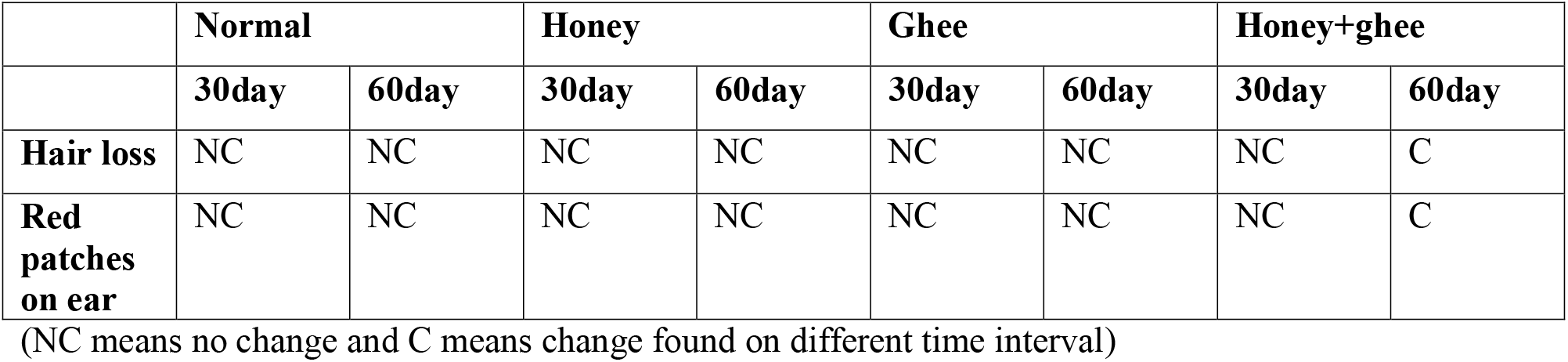
Physical appearance of different group’s rat on different time interval.

|  | Normal |  | Honey |  | Ghee |  | Honey+ghee |  |
| --- | --- | --- | --- | --- | --- | --- | --- | --- |
|  | 30day | 60day | 30day | 60day | 30day | 60day | 30day | 60day |
| **Hair loss** | NC | NC | NC | NC | NC | NC | NC | C |
| **Red patches on ear** | NC | NC | NC | NC | NC | NC | NC | C |

(NC means no change and C means change found on different time interval)

### 3.2. Effect on Biochemical parameter (Table:3)

#### 3.2.1. Liver function test

Plasma SGPT, SGOT, ALP, Protein, albumin, hemoglobin level was found similar in honey, ghee and the normal group. SGOT, SGPT, ALP, Protein and albumin level was found 7%, 24%, 25%, 9.8% and 25% increased respectively in the honey+ghee group in comparison to the normal group. Hemoglobin level was found 27% decreased in the honey+ghee group.

**Table 3:**
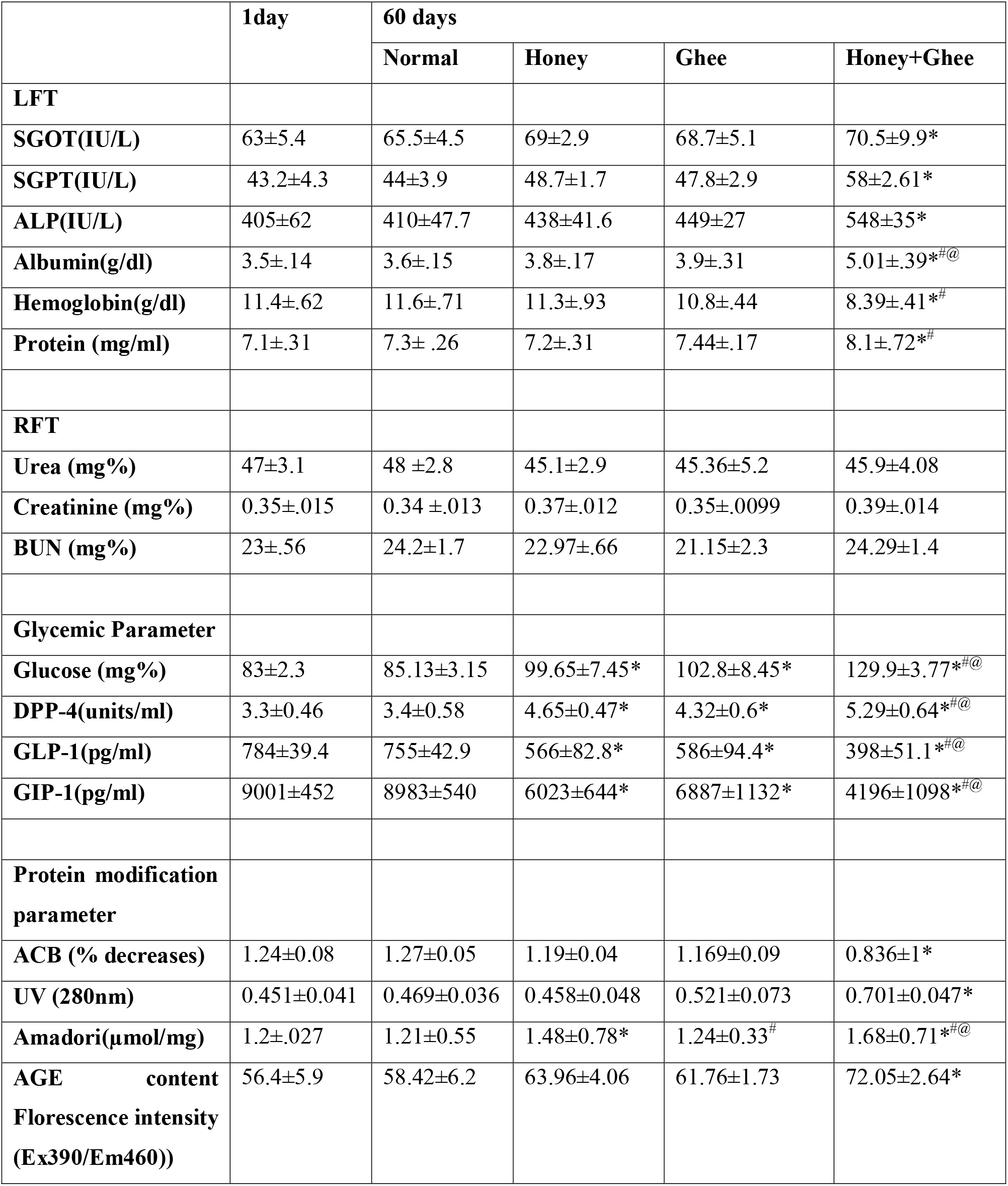
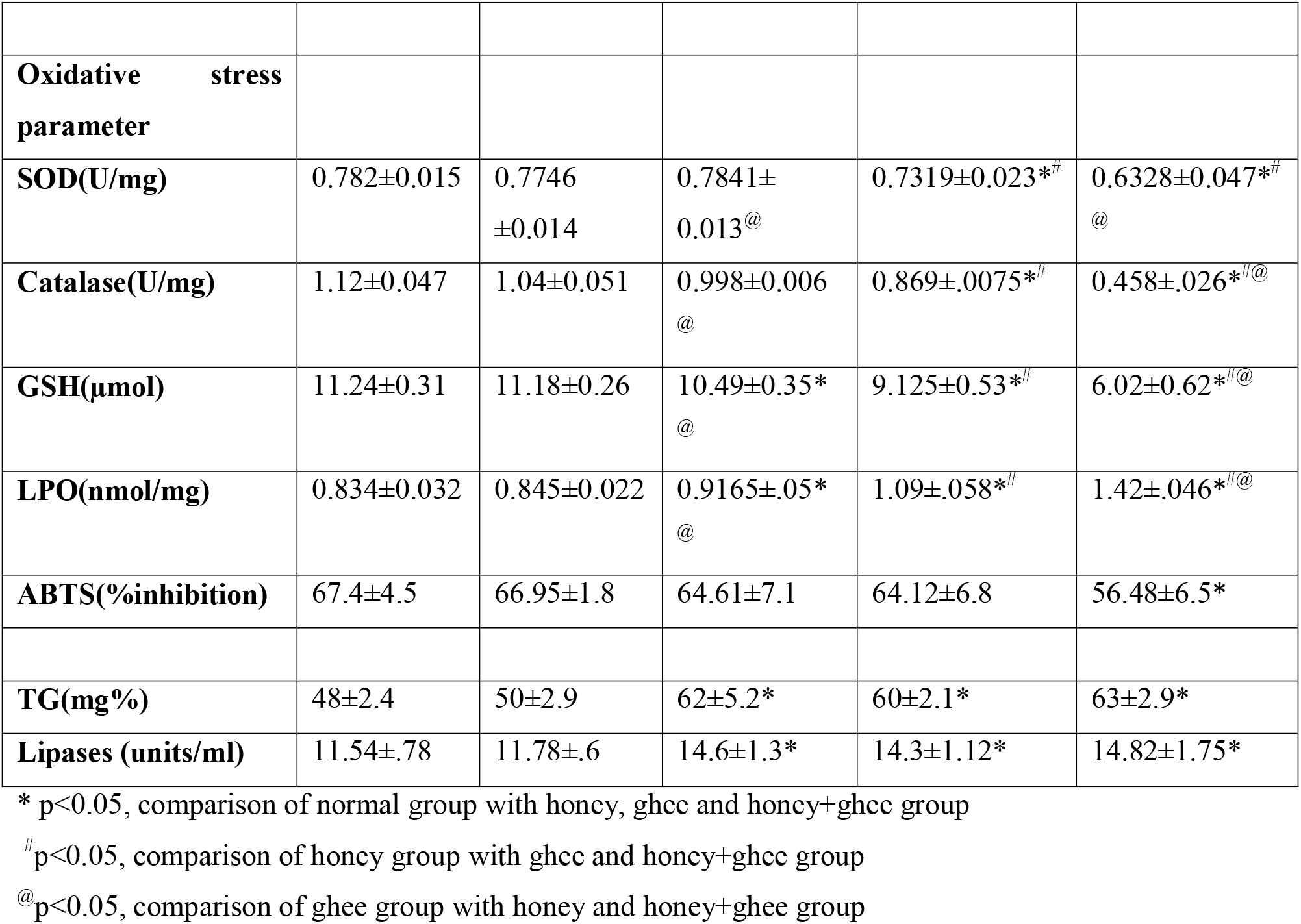
Effect of honey, ghee and equal ratio of honey and ghee on biochemical parameter.

#### 3.2.2. Kidney function test

Plasma urea, creatinine and blood urea nitrogen level was found similar in all groups.

#### 3.2.3. Oxidative stress parameter

Activity of blood superoxide dismutase and catalase was found 14% and 50% decreased respectively in honey+ghee group comparison to normal group. Level of reduced glutathione (GSH) was found 46% decreased in honey+ghee group. There was no any significant difference was found in honey group in comparison to normal. Level of lipid peroxidation was 40% increased in honey+ghee group in comparison to the normal group. Decreased % inhibition of ABTS^+^ indicated decreased free radical scavenging activity in the honey+ghee group.

#### 3.2.4. Glycemic parameter

Plasma glucose level was found 14%, 16% and 34% increased in honey, ghee and honey+ghee group respectively in comparison to the normal group. Plasma DPP-4 level was found 29%, 23% and 37% increased in honey, ghee and honey+ghee group respectively in comparison to the normal group. Plasma GLP-1 level was found 27%, 24%, and 49% decreased in honey, ghee and honey+ghee group respectively in comparison to the normal group. Plasma GIP-1 level was found 31%, 21% and 51% decreased in honey, ghee and honey+ghee groups respectively in comparison to the normal group.

#### 3.2.5. Protein modification parameter

Amadori product formation in plasma was found markedly increased 18% in honey and 27% in the honey+ghee group. UV absorption on 280 nm was 33% increased in the honey+ghee group plasma sample. Albumin cobalt binding capacity was found significantly decreased (34%) in honey+ghee group. Increased fluorescence intensity (18%) showed increased advance glycation ends product formation in honey+ghee group.

#### 3.2.6. Triglycerides and Lipases level

Triglycerides level was found increased 19, 16, and 20 % in honey, ghee and honey+ghee group respectively in comparison to the normal group. Lipases level was found increased 18.5 %, 17%, and 20 % in honey, ghee and honey+ghee group respectively in comparison to the normal group.

### 3.3. Histological study

H&E staining of liver tissue of normal, honey and ghee group rat showed normal morphology and no inflammation. Honey+ghee group rat liver tissue showed neutrophil infiltration, glycogen accumulation in some nucleus, some amount of bile duct dilation, and mild inflammation in portal tract which indicates initiation of inflammation in liver tissue (Figure 5). H&E staining of pancreas tissue of honey, ghee showed normal morphology with normal Islet of Langerhans but honey+ghee group showed dilated and thick blood vessel but no change in Islet of Langerhans (Figure 6). H&E staining of kidney tissue (Figure 3)showed normal morphology in all groups. H&E staining of intestine (jejunum) (Figure 4) tissue showed normal morphology in all four groups.

**Figure 3:**
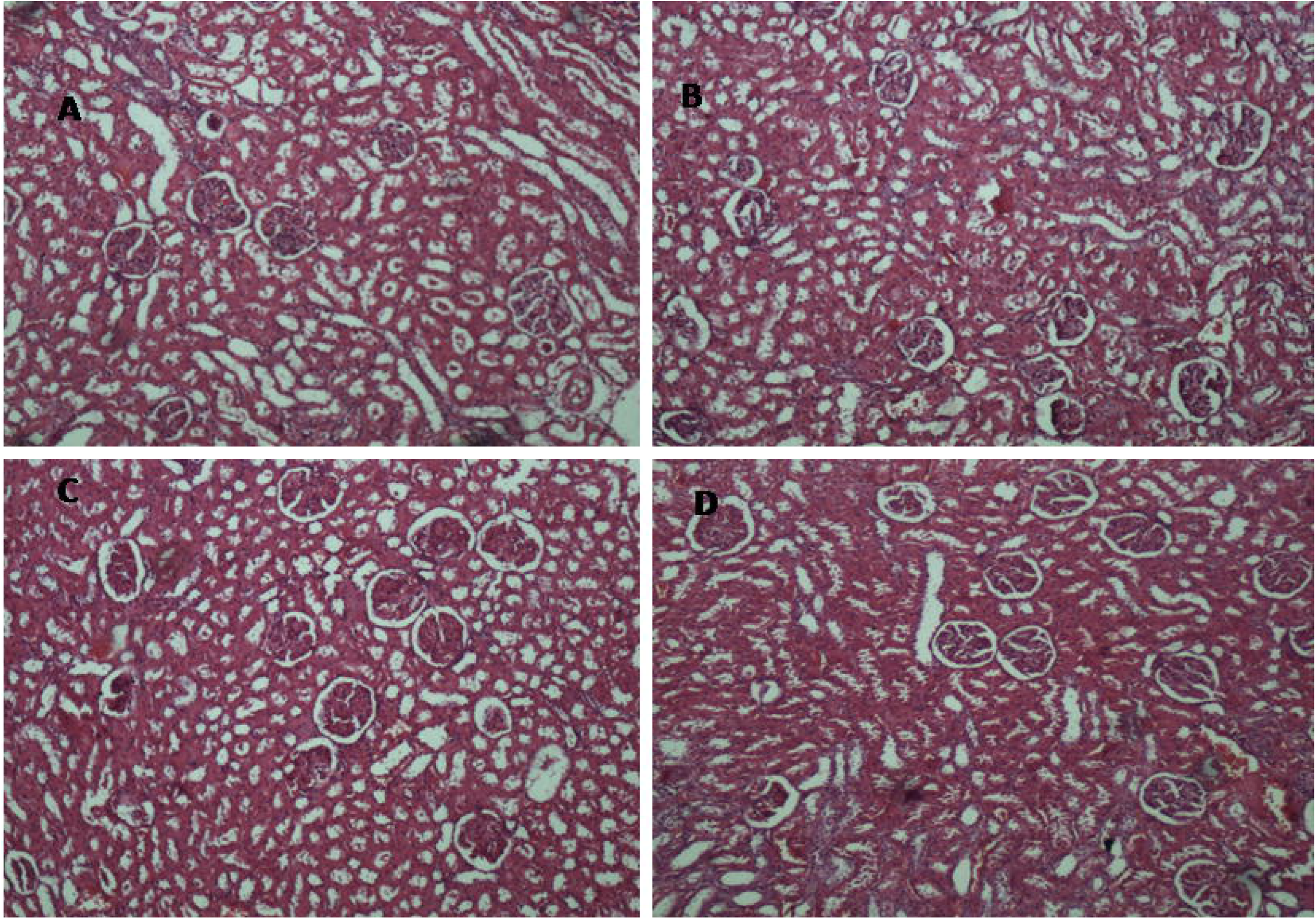
H&E image of Kidney tissue (10X), A (Normal), B (Honey), C (Ghee), D (Honey+Ghee). This image showed the normal morphology of kidney tissue in all groups.

**Figure 4:**
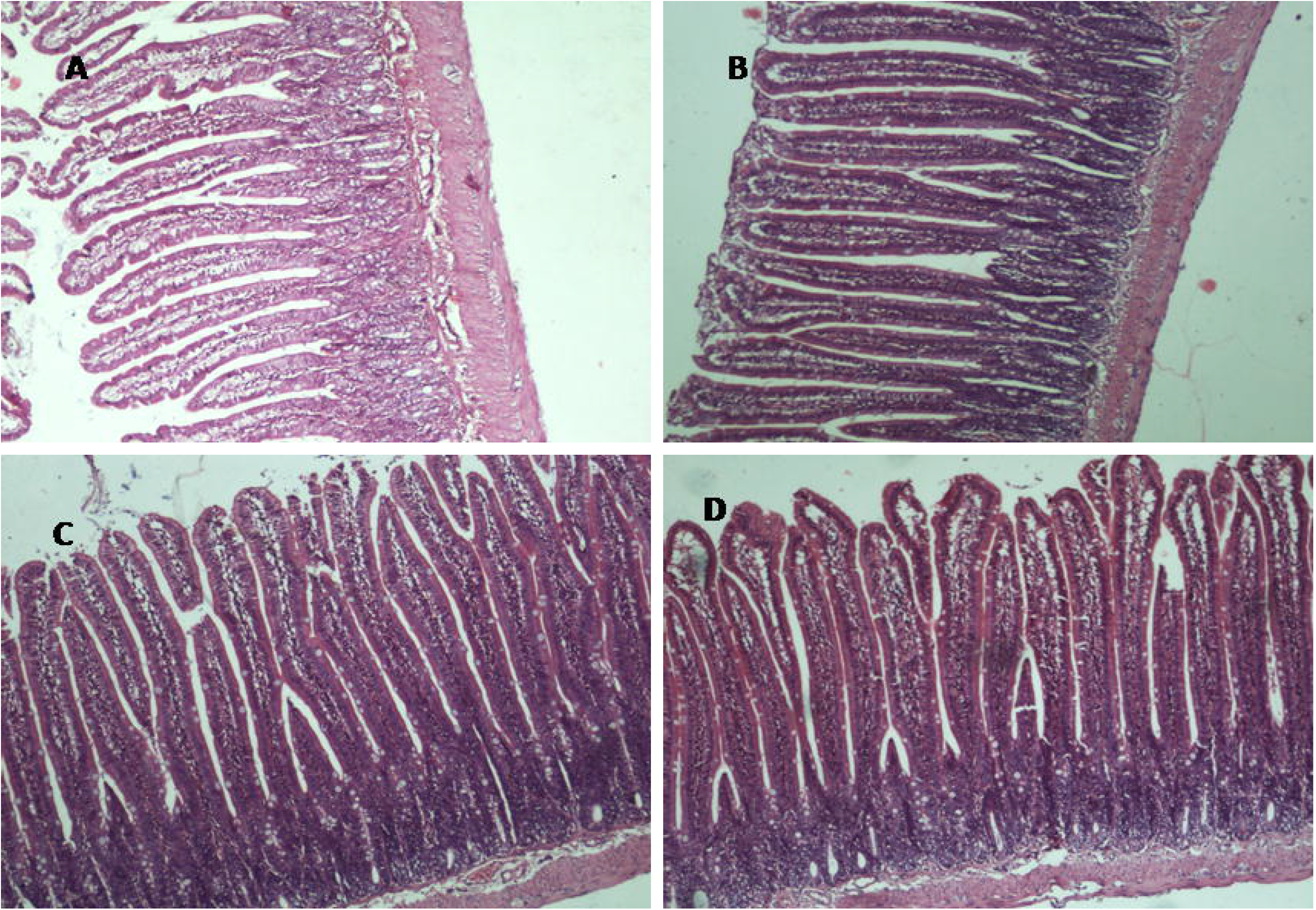
H&E image of intestine (jejunum) tissue (10X), A (Normal), B (Honey), C (Ghee), D (Honey+Ghee). This imaged showed the normal morphology with normal villi length in all four groups.

**Figure 5:**
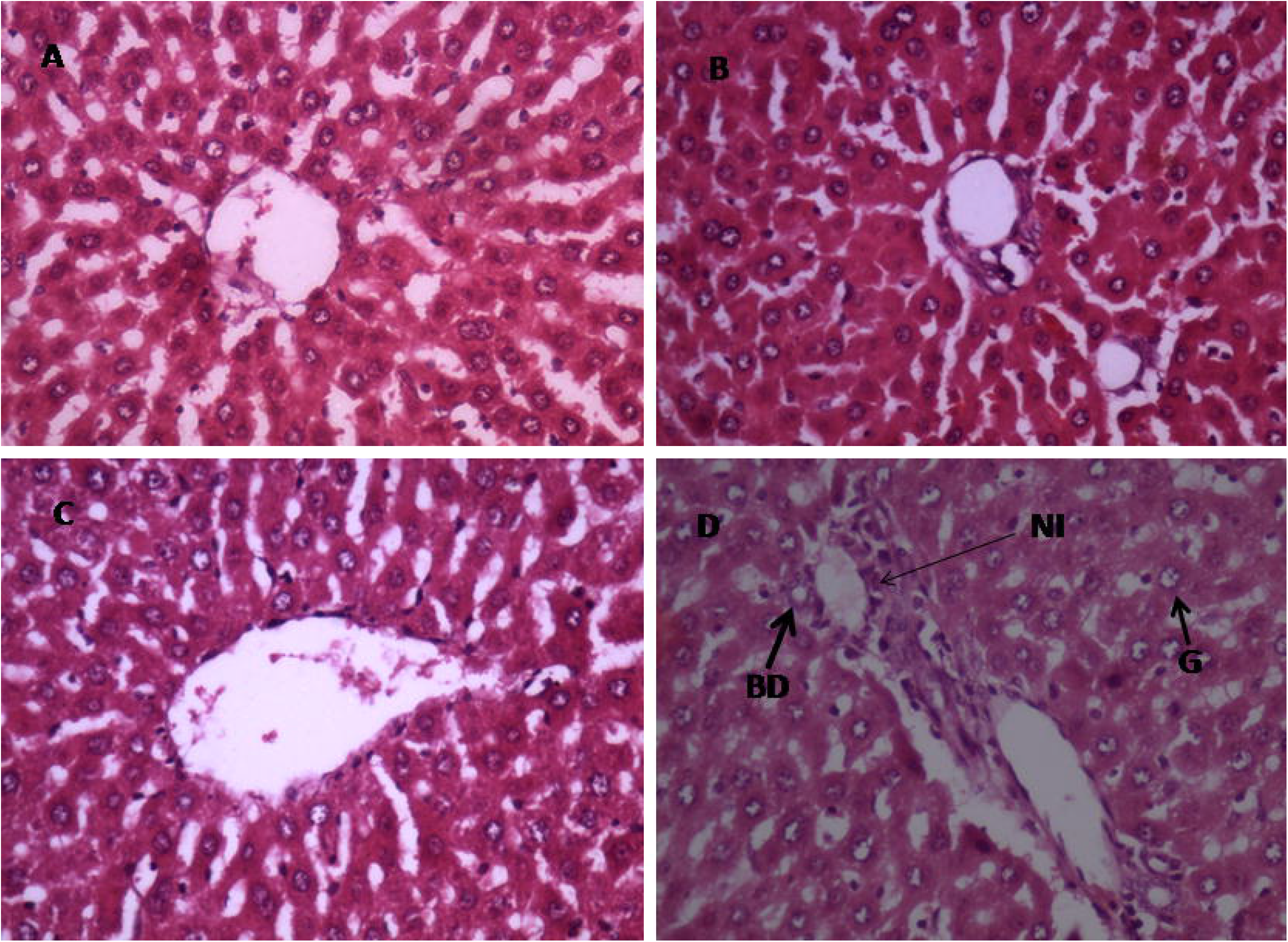
H&E image of liver tissue (40X). A (Normal), B (Honey), C(Ghee), D (Honey+Ghee). In honey+ghee showed some amount of accumulation of glycogen(G) in nucleus, neutrophil infiltrations(NI), bile duct dilation(BD) comparison than the normal, honey and ghee.

**Figure 6:**
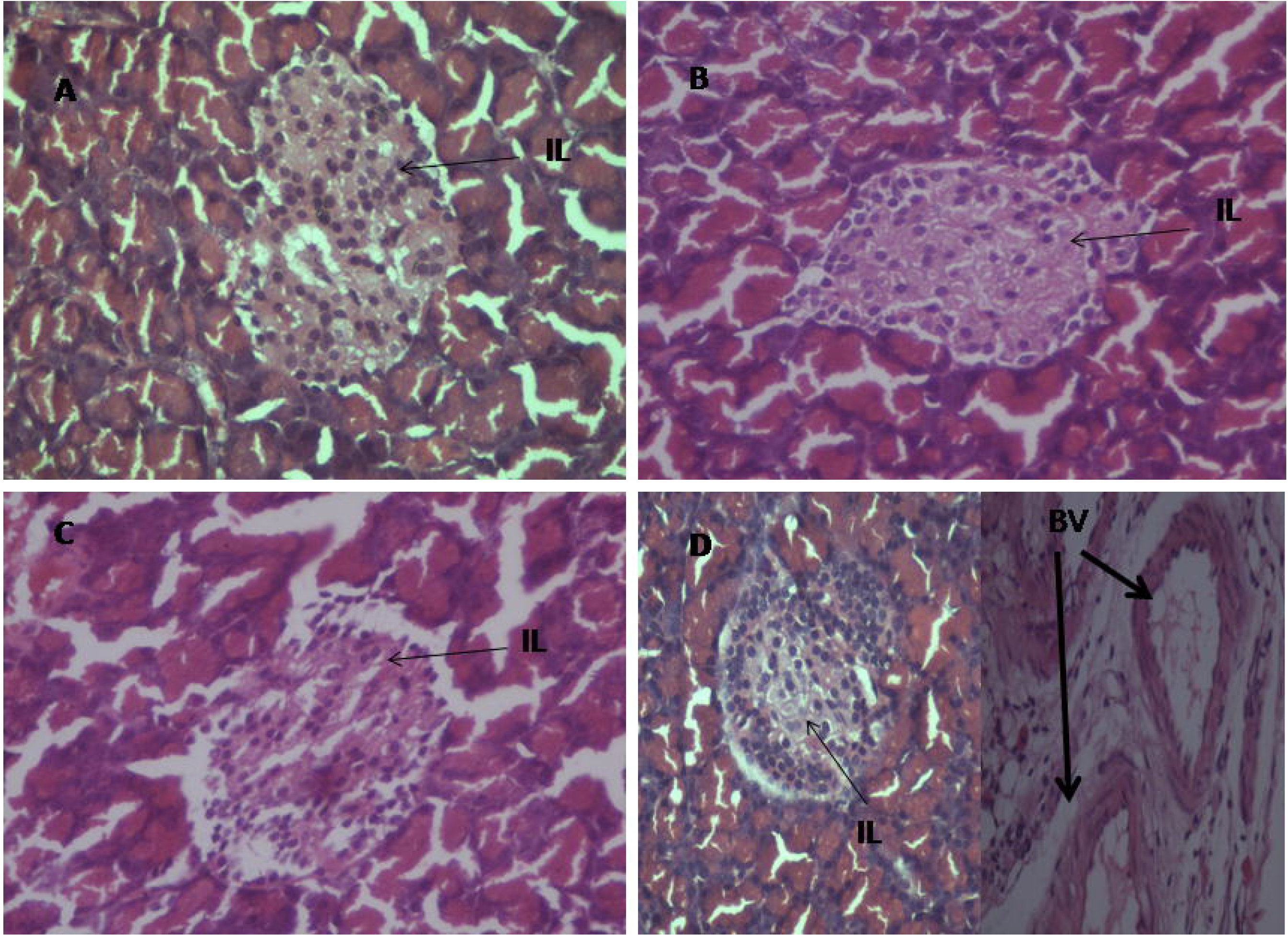
H&E image of Pancreas tissue (40X), A (Normal), B (Honey), C (Ghee), D (Honey+Ghee) (Arrow shows the normal islet of Langerhans (IL), in image D some level of dilated and thick blood vessel were found (BV, thick arraow))

## 4. Discussion

This present study has been focused on to know the reason behind an equal mixture of honey and ghee which is reported to be toxic in long used. In this study, weight of rats, treated with equal mixture of honey and ghee was significantly decreased in comparison to other groups. These rats also showed hair loss and skin patches. Such effects have been reported in patients going under chemotherapy, immune check point inhibitor, program protein inhibitor (PD1) at the latter stage of treatment even after discontinuing the medicine. In these patients the symptoms of Alopecia (hair loss), skin patches, dermatitis eczema are observed(Sharma et al., 2018). The genes like Tnfrs19, Ercc2, Lama5, Ctsl, Per1 are shown to be involved in etiology of skin disorders, these genes regulate various biological process involved in hair follicle development, hair follicle maturation, hair follicle morphogenesis and regulation of hair cycle(Lim, Kim, Lim, & Kim, 2019). Thus our observations showing the toxicity of honey and ghee are supported by other toxins reported here. The biochemical mechanism behind these toxicity could be associated with oxidative stress as it has been reported in case of hair loss and alopecia (Prie et al., 2015).

Literature showed that the AGEs is responsible for the glycation of many extracellular membrane protein like collage, elastin (Lim et al., 2019). Then the glycated extracellular protein is no longer responsible for the tightness of the skin, so the skin becomes aged and loses. This looseness of skin is responsible for the loss of hair and weight as well. This could be possible caused behind the hair loss and weight loss in rats of the honey+ghee group in our experiments.

In our study, we found a decreased level of SOD, catalase, GSH, % inhibition of ABTS and increased level of LPO. It is known that honey is rich in phenolic compound, flavanoids, vitamins and proteins which give honey antioxidant properties. As well as ghee also contain some vitamins which give antioxidative properties to ghee. But when we mix honey and ghee in equal ratio there is some chemical changes occur which is responsible for the formation of a carcinogenic compound hydroxyl methyl furfural which may produce the deleterious effect(K. Anilakumar et al., 2010). These hydroxyl methyl furfural is already present in honey in very little amount. The increased amount of this HMF production makes honey also toxic. The furfural and hydroxyl methyl furfural is formed by the acid catalyzed degradation of pentoses and hexoses respectively. The HMF formation is directly correlated with the chemical characteristic, free acids content, pH, metal content, lactone content and total acidity. This hydroxyl methyl furfural has found to be accountable for the induction and accumulation of reactive oxygen species (ROS), which is ultimately responsible for the cell component death in all organ including mitochondria and vacuolar membrane and chromatin in yeast (Allen et al., 2010).

The primary oxygen radical species generated by mitochondria is superoxide anion which is converted to hydrogen peroxide (H_2_O_2_) by spontaneous dismutation or by superoxide dismutase (SOD), present both within the mitochondria and in the cytosol. Hydrogen peroxide, in turn, is converted into water by glutathione peroxidase or catalase; otherwise, in the presence of divalent cations such as iron, H_2_O_2_ can undergo Fenton’s reaction to produce hydroxyl radical OH. Due to increased production of ROS, the activity of these enzymes like SOD, GSH, catalase are increased in the beginning to overcome the oxidative stress indicating the adaptive step but when ROS productions increases continuously then these antioxidative enzymes are unable to overcome oxidative stress and then level are decreases. One of the study showed that the furfural and HMF create oxidative stress by acting thiol-reactive electrophiles(Kim & Hahn, 2013), which relates to decreases GSH level.

In addition, this may block free ^34^SH of albumin and reducing its antioxidant properties. In this study, we found an increased level of albumin but decreased level of albumin cobalt binding which was due to oxidative stress. The free ^34^SH group of albumin is responsible for the binding of cobalt and other metal ions for transportation in the blood but when SH group is oxidized then the capacity of albumin to cobalt binding is decreased which is the sign of oxidative stress. The decrease in % inhibition of ABTS indicates the decreased free radical scavenging activity and increased lipid peroxidation also indicates the oxidative stress.

There is various type of oxidoreductase enzyme which carried out the reductive detoxification of furfural and HMF(Liu & Moon, 2009)(Lewis Liu, Moon, Andersh, Slininger, & Weber, 2008)(Almeida et al., 2008). During the reduction process, a lot of NADH and NADPH used as oxidoreductases cofactors. The consumption of NADH and NADPH for the reduction of furfural and HMF is responsible for the inhibition of different types of biosynthetic pathways which creates toxicity in the cell or tissues.

The amadori product formation occurs when the protein and glucose level is increase in blood which is the basis of the formation of advanced glycation end products (AGEs). Here in the present study, we found that the significant increase in amadori product formation and advanced glycation end product formation. The formation of amadori product leads to the formation as well accumulation of reactive carbonyl species through dehydration and rearrangement of amadori product. The major carbonyl intermediates are glyoxal, methylglyoxal and 3-deoxyglucosaone((Skovsted, Christensen, Breinholt, & Mortensen, 1998)(Baynes, 1991)). These compounds are responsible for intracellular oxidative stress which cause cellular apoptosis (Okado et al., 1996). AGEs have long been measured as poisonous that promote cell death and cause organ damage in man. AGEs is not responsible only for diabetic complication as well as it contributes for the development of many others metabolic diseases such as cardiovascular diseases (CVDs) and neurodegenerative diseases and aging(Van Puyvelde, Mets, Njemini, Beyer, & Bautmans, 2014).

We found in our study that the absorption of serum protein was notably increased in the group of equal mixture of honey and ghee fed rats than the other groups. These changes in absorption could be due to the formation of AGEs and oxidation of protein. Here we found decreased level of hemoglobin; this may be due to the glycation of hemoglobin. Increased amadori product formation may be due to the glycation of hemoglobin.

Honey is reported as a antidiabetic agent but when we take it in higher doses and for longer time the fructose and glucose present in it may responsible for the augmentation in glucose level because fructose is responsible for the creation of insulin resistant and obesity development(Matsushita, Ishii, Kijima, Jin, Takasu, Kuroda, et al., 2012). Long term carbohydrate and fat rich diet feeding is linked to the hyperglycemia condition. Increased glucose level is responsible for activation of DPP-4 enzyme. DPP-4 is an intestinal enzyme responsible for inactivation of GLP-1 because the sustained activity of GLP-1 is not recommended for normal physiology. The normal life for GLP-1 is 3-4 minute thus the optimum activity of DPP-4 is desired for a healthy person but increases in DPP-4 is injurious to health because it damages the normal level of GLP-1 and GIP-1 also(Mentlein, Gallwitz, & Schmidt, 1993). Glucagon-like peptide-1 (GLP-1) is a remarkable antidiabetic gut hormone with combinatorial actions of stimulating insulin secretion, inhibiting glucagon secretion, increasing beta-cell mass, reducing the rate of gastric emptying and inducing satiety. Gastric inhibitory polypeptide-1(GIP-1) inhibits gastric acid secretion and stimulates insulin secretion in healthy volunteers. 3-deoxyglucosone is an amadori product. Accumulation of 3-deoxyglucosone in intestinal tissue is responsible for the attenuation of GLP-1 secretion which causes insulinotropic effect in rats(Zhang et al., 2016). For the first time we are reporting the changes in incretins hormones by using equal amount of honey and ghee which are not reported earlier. Further study are required this linked to toxicity.

The increased formation of amadori product and advanced glycation end product leads to direct inflammation in liver by various types of signaling pathways((Hyogo & Yamagishi, 2008)(Bijnen et al., 2019)). HMF is also responsible for the mutation of important gene in liver(Matsushita, Ishii, Kijima, Jin, Takasu, & Kuroda, 2012). The possible cause of liver inflammation in honey+ghee group rats could be the formation of these HMF, amadori product, advanced glycation end product and oxidative stress. We found that the liver function test enzymes serum glutamate oxaloacetate transaminases (SGOT), serum glutamate pyruvate transaminases (SGPT), alkaline phosphatases(ALP) activity were significantly increased in honey+ghee group in comparison to others group. These enzymes are the initial marker of liver inflammation. The increased amount of protein, albumin in honey+ghee group in comparison to other three groups also indicates the early sign of liver inflammation. H&E image of liver tissue showed mild inflammation in portal track, neutrophil infiltration, some amount of glycogen accumulation in the nucleus and some bile duct dilation which showed the initiation of inflammation in equal mixture of honey and ghee treated rats.

In this study, we didn’t find any significant differences in blood urea, Creatinine, blood urea nitrogen and kidney histology between all groups, which indicates that the equal mixture of honey and ghee has no effect on kidney till 60 day.

When we take carbohydrate-rich diet it is directly related to fatty acid synthesis and further triglycerides. A fat rich diet contains many fats, triglycerides and cholesterol which are directly linked to the TG accumulation in liver and blood also. Enzyme lipase is responsible to the degradation of these triglycerides into fatty acids. We found the similar result in our experiments.

## 5. Conclusion

As discussed above, increased oxidative stress generation, decreased albumin cobalt binding, increased amadori product formation and advance glycation end product formation, glucose and DPP-4 augmentations which relate to GLP-1 and GIP-1 attenuation, liver function test enzyme elevation, liver tissue inflammation, neutrophil infiltration, bile duct dilation, glycogen accumulation, rise in TG and lipases level are the evidence which can say that the formation of HMF, amadori product like 3-deoxyglucosone, advance glycation end product could be the possible cause of toxicity of equal ratio of honey and ghee. These may be responsible for weight and hair loss as well as the appearance of red patches on ears.

## Abbreviations

LFT: Liver function test,
RFT: Renal function test,
SGOT: Serum glutamate oxaloacetate transaminases,
SGPT: Serum glutamate pyruvate transaminases,
Alp: Alkaline phosphatases,
DPP-4: Dipeptidyl protease,
GLP-1: Glucagon-like peptide-1,
GIP-1: Gastric inhibitory polypeptide,
GPPN: gly-pro-p-nitroanilide,
ABTS: 2,2’-azino-bis-3-ethylbenzthiazoline-6-sulphonic acid,
HMF: Hydeoxymethyl furfural,
ACB: Albumin cobalt binding,
AGEs: Advanced glycation end products,
SOD: superoxide dismutases,
LPO: lipid peroxidation,
GSH: reduced glutathione,
HB: heamoglobin,
TG: triglycerides

## Acknowledgement

We would like to say thanks Prof. Mohan Kumar Sir for analyzing the H&E image of tissue sample of this study. We would like to say thanks to BHU-RET fellowship for funding me for this work.

