## Supplementary 1 for "Nutritive diet becomes unhealthy if taken in a wrong ratio: Scientific validation of Ayurvedic concept of “incompatible diet” (*Virrudh Aahar*)"

**Table 1: Nutrient information of honey and ghee (per 100 gm)**

|  | Honey (Patanjali) | Ghee (Anik) |
| --- | --- | --- |
| Energy | 320 kcal | 879 kcal |
| Carbohydrate | 80 g | 0.0g |
| Natural sugar | 80 g | 0.0g |
| Added sugar | 0 g |  |
| Protein | 1 g | 0.0g |
| Fat | 0 g | - |
| Sodium | 20 mg | - |
| Potassium | 130 mg | - |
| Calcium | 12 mg | - |
| Phosphorus | 5 mg | - |
| Iron | 1.6 mg | - |
| Water | - | 0.3g |
| Fibre | - | 0.0g |
| Cholesterol | - | 275-325mg |
| Saturated fatty acids | - | 58g |
| Monounsaturated fatty acids | - | 28g |
| Polyunsaturated fatty acids | - | 2g |
| Trans fatty acids | - | 5g |
| Vitamin A | - | 2000-3500IU |

% CHANGES

|  | SOD | Catalase | GSH | LPO |
| --- | --- | --- | --- | --- |
| 1 DAYS | 0 | 0 | 0 | 0 |
| 15 DAYS | 5.74 | 15.46 | 9.9 | 1.3 |
| 30 DAYS | -9.41 | -32.54 | -44.8 | 16.34 |
| 45 DAYS | -11.62 | -126.2 | -48.35 | 29.32 |
| 60 DAYS | -16.45 | -144.2 | -88.96 | 40.23 |

Figure: % change in oxidative stress parameter on different days of honey+ghee group.
